# Antisense Oligonucleotides Targeting an *LDLR* Regulatory RNA Increase *LDLR* Expression and Reduce LDL-cholesterol in Vivo

**DOI:** 10.64898/2026.06.02.729508

**Authors:** Keerthana Gnanapradeepan, Salomé Manska, Brian Schwartz, Gabriel Golczer, Jiaqi Huang, Rachana S. Kelkar, Andrew Masteller, Shama Pilankar, Marianne Richter, Scott Waldron, Brynn N. Akerberg, Dale C. Guenther, Alla A. Sigova, David Bumcrot, Daniel F. Tardiff

**Affiliations:** CAMP4 Therapeutics Corp., Cambridge, MA USA

## Abstract

Regulatory RNAs (regRNAs) are non-coding RNAs expressed from promoters and enhancers that modulate gene transcription. Targeting regRNAs with antisense oligonucleotides (ASOs) can increase transcription and presents a potential therapeutic strategy to restore expression in haploinsufficient diseases. This approach requires high-quality regRNA maps, identification of ASOs targeting actionable sequences, and translation of efficacy to relevant animal models. Here, we describe this approach for human low-density lipoprotein receptor (*LDLR*). We characterized *LDLR* regRNAs in HepG2 cells and human liver using epigenomic mapping and regRNA Capture-seq, and designed ASOs targeting these regRNAs. Several ASOs increased *LDLR* expression in cells and a lead was evaluated for efficacy in a humanized liver mouse model. ASO administration increased human *LDLR* mRNA and protein, while lowering LDL-cholesterol in plasma. These data provide a framework for the discovery and translation of ASOs that increase gene expression as a potential therapeutic approach for the treatment of haploinsufficient diseases.

## Introduction

RNA therapeutics that exploit hybridization-dependent mechanisms to selectively target RNAs and modulate their fate have the potential to address a large number of human diseases. Both small interfering RNAs (siRNA) and antisense oligonucleotides (ASO) are widely used to selectively reduce levels of target mRNAs with the intent of ameliorating underlying causes of disease. ASOs that recruit RNase H1 to specifically cleave their cognate mRNA result in mRNA knockdown and subsequent protein reduction^1^. However, many diseases, such as those caused by insufficient protein levels, or haploinsufficiency, will not benefit from a knockdown mechanism. Approaches to increase protein levels, although less explored, hold significant potential to expand the scope of oligonucleotide applications. For example, splice-switching oligonucleotides (SSO) have the potential to boost protein levels in diseases such as Spinal Muscular Atrophy, Duchenne’s muscular dystrophy, and Dravet Syndrome^2–4^. Other ASO-mediated mechanisms, such as stabilizing mRNA, inhibiting translation of upstream open reading frames, or targeting natural antisense transcripts, have the potential to modulate mRNA function and increase protein levels^5–8^.

An untapped class of RNAs that are becoming increasingly appreciated as key regulators of transcription are non-coding regulatory RNAs (regRNAs), which are expressed from both enhancers (eRNAs) and promoters (paRNAs)^9,10^. Transcription factors (TFs) bind these RNAs and such interactions help maintain high local concentrations of TFs, likely within transcriptional condensates, to regulate gene transcription^11,12^. Epigenomic characteristics, including histone modifications, chromatin accessibility, and three-dimensional chromatin organization, can be integrated with measures of active transcription to identify regulatory regions^13,14^. A recently described method, called regRNA Capture-seq, was developed to isolate and sequence low-abundance RNAs expressed from regulatory regions in in primary human hepatocytes^15^. Importantly, in addition to generating this large catalog of regRNAs, the role of an ornithine trans-carbamylase (*OTC*) regRNA in modulating transcription was elucidated through the identification of ASOs that specifically targeted the regRNA. These ASOs increased transcription by a mechanism that required regRNA structure perturbation and altered transcription factor binding^15^.

Here, we describe a framework for the characterization of regRNAs, discovery of ASOs resulting in gene upregulation, and validation in a relevant human model using the *LDLR* gene. LDLR is a receptor for low density lipoprotein-cholesterol (LDL-c) expressed on the surface of hepatocytes and is mutated in heterozygous familial hypercholesterolemia (HeFH), resulting in LDLR haploinsufficiency and elevated LDL-c^16^. We used both epigenomic mapping and regRNA Capture-seq to characterize the *LDLR* regRNA landscape, which enabled the identification of ASOs that increased *LDLR* expression in vitro. Critically, targeting the *LDLR* regRNA also enhanced gene expression and reduced LDL-c in a humanized liver mouse model. These data illustrate how regRNAs can be characterized and targeted with ASOs for the potential treatment of haploinsufficient diseases.

## Results

### Characterization of *LDLR* regRNA landscape

*LDLR* was selected as a model target gene to both identify regRNAs using advanced genomic methods and to discover ASOs that increase gene expression. The availability of a soluble biomarker in LDL-c provided a translational path to proof of concept, and the prior initial description of a paRNA expressed antisense from the promoter provided a comparison to our approach^17,18^. We first mapped promoter-proximal regulatory elements at the *LDLR* locus in HepG2 cells and human liver tissue. Our analysis focused on a ∼3.5 kb region surrounding the annotated *LDLR* transcription start site (TSS). Candidate promoters and enhancers were identified by integrating assay for transposase-accessible chromatin sequencing (ATAC-seq) with chromatin immunoprecipitation sequencing (ChIP-seq) profiles for histone H3 lysine 4 trimethylation (H3K4me3), H3 lysine 4 monomethylation (H3K4me1), and H3 lysine 27 acetylation (H3K27ac). Precision Run-On sequencing (PRO-seq) was used to assess transcriptional activity across the locus (Fig. 1A, Supplementary Fig. 1A).

**Fig. 1.**
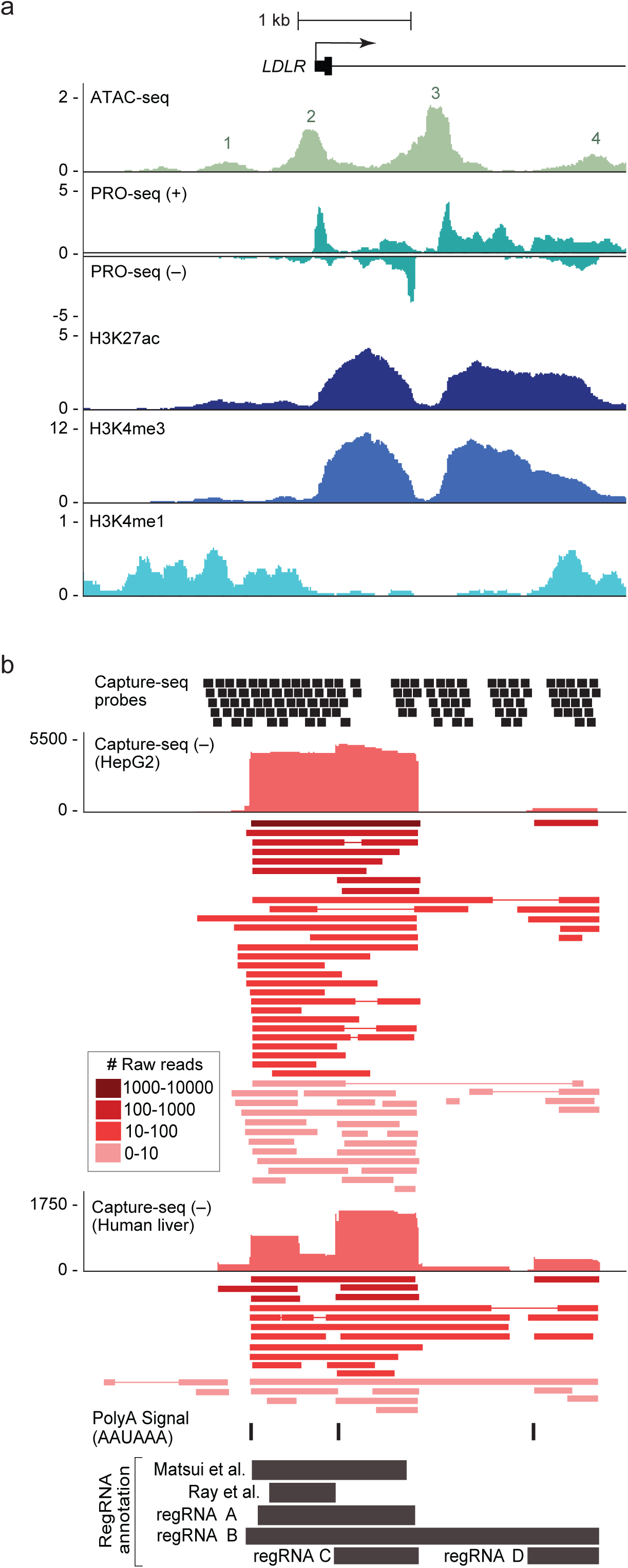
Mapping of chromatin interactions and identification of regulatory RNAs at the *LDLR* locus. A) Signal tracks at the *LDLR* locus in HepG2 cells. ATAC-seq (light green) and PRO-seq (teal) peaks are represented as reads per million (RPM). ATAC-seq summits are numbered 1-4. PRO-seq sense (+) and PRO-seq anti-sense (–) reads are displayed above and below the y-axis, respectively. ChIP-seq signals for H3K27ac (dark blue), H3K4me3 (medium blue), and H3K4me1 (light blue) are represented as RPM. B) Capture-seq signal tracks at the *LDLR* locus in HepG2 cells and human liver showing captured regRNA species. Probe design tiling for Capture-seq are displayed with black bars representing a single 80nt biotinylated RNA probe for hybridization. Negative-strand (−) Capture-seq regRNA reads (red) are shown as raw read peak intensity with each captured regRNA species below. Raw read abundance for captured regRNA species is indicated by the dark-to-light color scale in the legend. PolyA motifs (AAUAAA) are indicated, and annotated regRNAs are displayed below the tracks.

Two prominent ATAC-seq peaks separated by ∼1 kb (Fig. 1A; peaks 2 & 3) marked accessible chromatin and overlapped with PRO-seq signal, indicating convergent transcription initiation at the *LDLR* TSS in the sense direction and from the downstream ATAC-seq peak in the antisense direction (Fig. 1A). Enrichment of H3K4me3 and H3K27ac at these loci confirmed that both correspond to active promoter elements (Fig. 1A). In contrast, the weaker ATAC-seq peaks (Fig. 1A; peaks 1 and 4) were associated with H3K4me1-modified nucleosomes, indicative of enhancer regions. Of these, only the downstream enhancer also displayed H3K27ac-modified nucleosomes and detectable PRO-seq signal, consistent with an active state, whereas the upstream enhancer lacked both signals, consistent with a poised or inactive element. Together, these data indicate the presence of two promoters and one active enhancer within the proximal *LDLR* regulatory landscape.

To comprehensively characterize polyadenylated transcripts arising from these regulatory elements, we designed regRNA Capture-seq probes targeting all four regions of accessible chromatin (Fig. 1B). RegRNA Capture-seq was selected for its strand specificity, higher sensitivity for low-abundance transcripts, and superior positional resolution relative to other RNA detection approaches. By using long-read sequencing, this method enables precise mapping of transcript boundaries, isoform complexity, and relative abundance^15^. RegRNA Capture-seq identified multiple partially overlapping RNA species transcribed from both the sense and antisense DNA strands in HepG2 cells and human liver tissue. Sense transcripts originating at the annotated *LDLR* TSS corresponded to the alternatively spliced *LDLR* mRNA isoforms with variable 3′ untranslated region (UTR) lengths (Supplementary Fig. 1). In contrast, multiple antisense-strand transcripts originated from both *LDLR* promoters and the active enhancer (Fig. 1B; Supplementary Fig. 1).

These promoter- and enhancer-associated regRNAs - annotated as regRNAs A, B, C, and D - were non-coding, predominantly unspliced, substantially shorter than *LDLR* mRNA, and ranged from ∼100 to 3,000 nt in length (Fig. 1B). All major regRNAs contained canonical polyadenylation signals near their 3′ ends and shared partial sequence homology with multiple shorter species exhibiting heterogenous 3’ and 5’ends, consistent with cryptic polyadenylation and potential alternative transcription initiation. Notably, a subset of regRNAs displayed alternative splicing (regRNAs A and D). In addition to novel spliced and unspliced regRNA species (regRNAs B, C, D), we noted that unspliced regRNA A largely overlaps previously described antisense noncoding RNAs at the *LDLR* promoter^17,18^. Although regRNAs A–D were detected in both HepG2 cells and human liver tissue, their relative abundances differed between in vitro and in vivo contexts: regRNA A was more abundant in HepG2 cells, whereas a shorter ∼700 nt regRNA C was more abundant in liver tissue (Fig. 1B). Collectively, these findings reveal a complex and conserved population of regRNAs expressed in close proximity to the *LDLR* TSS, consistent with a potential role in the regulation of *LDLR* gene expression.

### Identification of ASOs targeting *LDLR* regRNA

Recent work has shown that targeting the *OTC* eRNA with ASOs can increase gene transcription^15^. With the *LDLR* regRNAs characterized, we asked whether ASOs complementary to the most abundant regRNA expressed in HepG2 cells, regRNA A, could increase *LDLR* gene expression. A set of 88 ASOs targeting *LDLR* regRNA A were individually transfected into HepG2 cells and *LDLR* gene expression was evaluated 48 hours later (Fig. 2A, Supplementary Fig. 2A-B, Supplementary Table 2). ASOs were designed with no assumptions about mechanism of action to allow for unbiased identification of potential hits. All ASO sequences were screened across two chemistries, one that supports steric binding without the possibility of cleavage (sterics) and the other that supports RNase H1-mediated cleavage (gapmer). These oligos were 20-nt in length with fully phosphorothioated backbones. Steric ASOs were fully modified with 2′-methoxyethyl (MOE) ribose modifications, whereas gapmer ASOs followed a 5-10-5 design, in which the internal 10 nucleotides contained unmodified DNA (2′-deoxyribose) to support RNase H1-mediated cleavage of the target RNA.

**Fig. 2.**
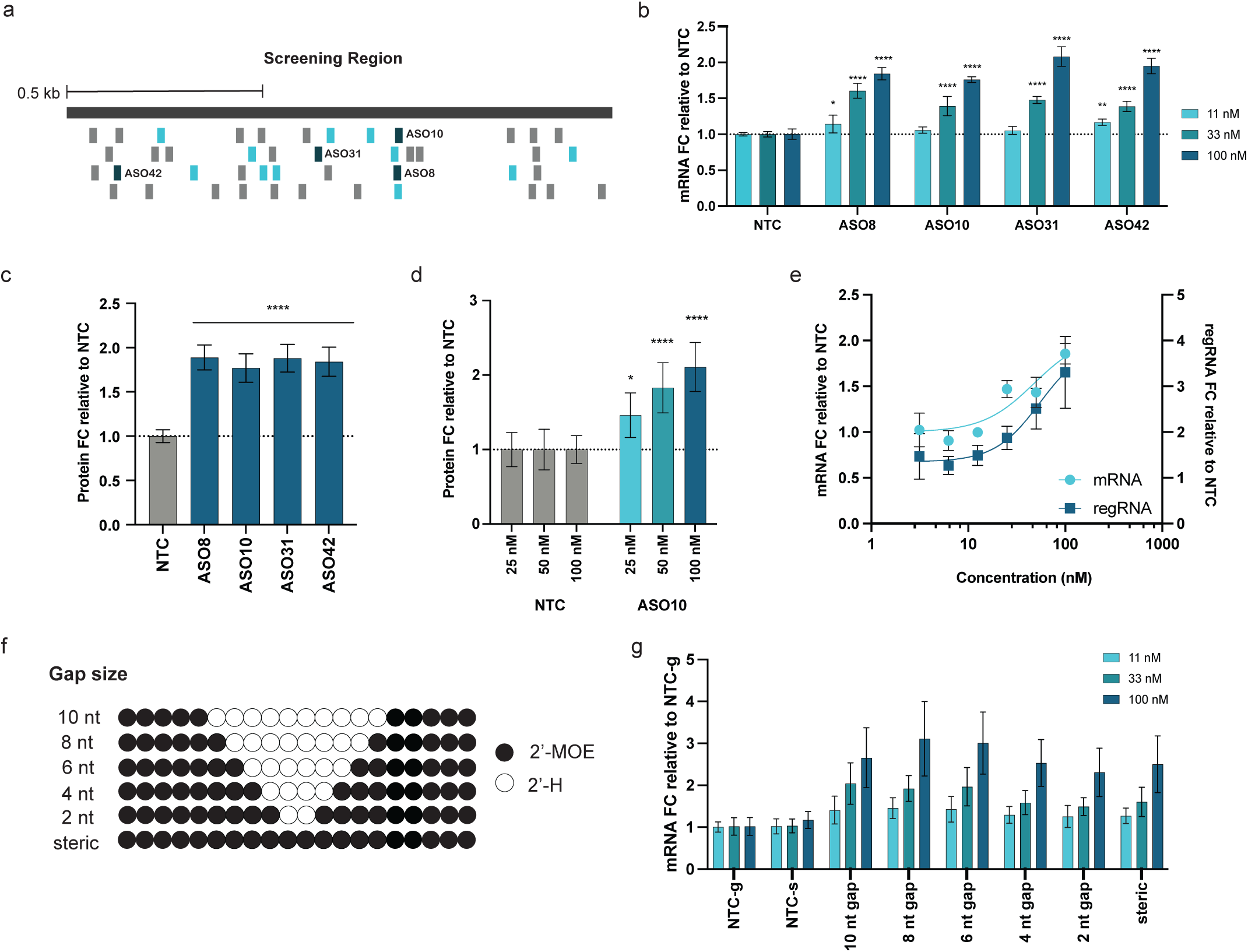
Characterization of ASO10 upregulation of *LDLR* in HepG2 cells. A) Location of screened ASOs targeting the *LDLR* regRNA screening region. Initial 15 hits identified from screen resulting in > 1.5x FC are shown in light blue; 4 hits prioritized for further characterization highlighted in dark blue and labeled. B) Validation of four prioritized primary screen hits. HepG2 cells were transfected with 11, 33 or 100 nM of ASO for 48 hours and assessed for *LDLR* expression using qRT-PCR. Data are presented as mean fold change relative to respective NTC concentration ± s.d. (n = 3; two-way ANOVA; ** p < 0.01, **** p < 0.0001). C) LDLR protein levels following treatment with prioritized screen hits. HepG2 cells were transfected with 100 nM ASO for 48 hours and LDLR protein was assessed using ELISA. Data normalized to total protein and shown as mean fold change relative to the NTC ± s.d. (n = 3; unpaired t test; **** p < 0.0001). D) LDLR protein concentration response for ASO10. NTC ASO or ASO10 were transfected at 25, 50, and 100 nM in HepG2 cells for 48 hours and LDLR protein was assessed using ELISA. Data were normalized to total protein and shown relative to the NTC ASO at corresponding concentrations ± s.d. (n = 3; two-way ANOVA; * p < 0.05, **** p < 0.0001). E) *LDLR* mRNA and regRNA concentration response curve for ASO10. NTC ASO or ASO10 were transfected at concentrations ranging from 3.1 to 100 nM into HepG2 cells for 48 hours and assessed for *LDLR* mRNA and regRNA expression using qRT-PCR. Concentration response curves were generated using a 4PL nonlinear regression model. Data are presented as mean fold change relative to respective NTC concentration ± s.d. (n = 3). *LDLR* mRNA is on the left y-axis; *LDLR* regRNA is on the right y-axis. F) Schematic depicting ASO10 that has a 5-10-5 gapmer design and derivatives with shortening DNA gap length. Black circles represent 2′-*O*-methoxyethyl, white circles represent unmodified DNA. G) *LDLR* mRNA in response to ASO10 derivatives with various DNA gap lengths. HepG2 cells were transfected with 11, 33 or 100 nM of ASO for 48 hours and assessed for *LDLR* expression using qRT-PCR to assess parent ASO and derivatives shown in D. Gapmer ASOs (10 nt to 2 nt) are normalized to NTC-g, steric ASO normalized to NTC-s. Data are presented as mean fold change relative to respective NTC ± s.d. (n = 3).

The majority of *LDLR*-regRNA A targeting ASOs had minimal effects on *LDLR* expression (Fig. 2 and Supplementary Fig. 2A-B). Fifteen ASOs (14 gapmers and 1 steric) were identified that increased *LDLR* mRNA by at least 1.5-fold relative to a non-targeting control (NTC) ASO with matched chemistry (Supplementary Fig. 2A-B). These fifteen ASOs were assessed in a three-point dose response, and ASO8, ASO10, ASO31 and ASO42 (all gapmers) demonstrated the most robust concentration-dependent upregulation with maximum increase of ∼2-fold at 100 nM (Fig. 2B, Supplementary Fig. 2C). Importantly, the increases in mRNA resulted in comparable ∼1.8-fold increases in LDLR protein measured by ELISA (Fig. 2C). ASO10, which partially overlaps with ASO8, was selected as a representative ASO for further studies based on its consistent upregulation of *LDLR* mRNA and protein. LDLR protein levels increased in a concentration-dependent manner with a maximal 2.1-fold increase at 100 nM and a 1.4-fold increase at the lowest concentration tested of 25 nM (Fig. 2D). Taken together, these data demonstrate that targeting the *LDLR* regRNA with ASO10 can increase both *LDLR* mRNA and protein in HepG2 cells.

### ASO10 increases *LDLR* through an RNase H1-independent mechanism

The gapmer structure of ASO10 supports the potential to direct RNase H1-mediated cleavage and reduce *LDLR* paRNA levels. We therefore assessed both *LDLR* mRNA and regRNA in a six-point concentration response curve of ASO10 (Fig. 2E). Despite its gapmer design, ASO treatment increased both *LDLR* regRNA and mRNA levels with comparable potencies of 54.5 and 47.6 nM, respectively. This result is consistent with prior studies that showed ASOs targeting regRNAs may increase not just expression of the target gene, but also the levels of the targeted regRNA^15^.

The observed increase in regRNA suggests that ASO10 may not be working through a cleavage mechanism, despite having a conventional gapmer design. To address this directly, derivatives in which the DNA gap length was reduced from 10 to 2 nucleotides, as well as a fully steric ASO, were synthesized and assessed for activity. All derivatives maintained comparable efficacy to ASO10 with maximal increases ranging from 2.3- to 3.1-fold (Fig. 2F-G). These data suggest that cleavage and reduction of the *LDLR* regRNA is not required for ASO10-mediated increases in *LDLR* gene expression. Rather it is likely that direct binding to the regRNA results in changes in regRNA function that mediate the observed increase in expression. Of note, ASO54 – the steric analog of ASO10 – had a 1.3-fold increase in the primary screen and was not selected for follow up based on the greater fold change of the gapmer ASOs (Supplementary Fig. 2). Performance of the steric was more robust when tested in concentration response experiments (Fig. 2G).

### GalNAc-conjugated ASO10 increases LDLR protein and reduces LDL-cholesterol in humanized liver mice

We next asked if ASO10 activity, and therefore the role of the *LDLR* paRNA in regulating expression, translated from cells to animals by assessing efficacy in a humanized liver mouse model (PXB Mouse^®^). These mice have livers repopulated with primary human hepatocytes and therefore express human *LDLR* and associated regRNAs, while also exhibiting overall lipid profiles more comparable to that of humans than mice (high LDL-cholesterol and low HDL-cholesterol)^19,20^.

ASO10 was conjugated to N-acetylgalactosamine (ASO10-Gal) to facilitate asialoglycoprotein receptor (ASGPR)-mediated uptake in hepatocytes. ASO10-Gal was then administered subcutaneously at 3 or 10 mg/kg on Days 1 and 15 (Fig. 3A). Blood was assessed for LDL-c and HDL-c, while livers were analyzed for *LDLR* mRNA, regRNA, protein and ASO concentration at the termination of the study on Day 29.

**Fig. 3.**
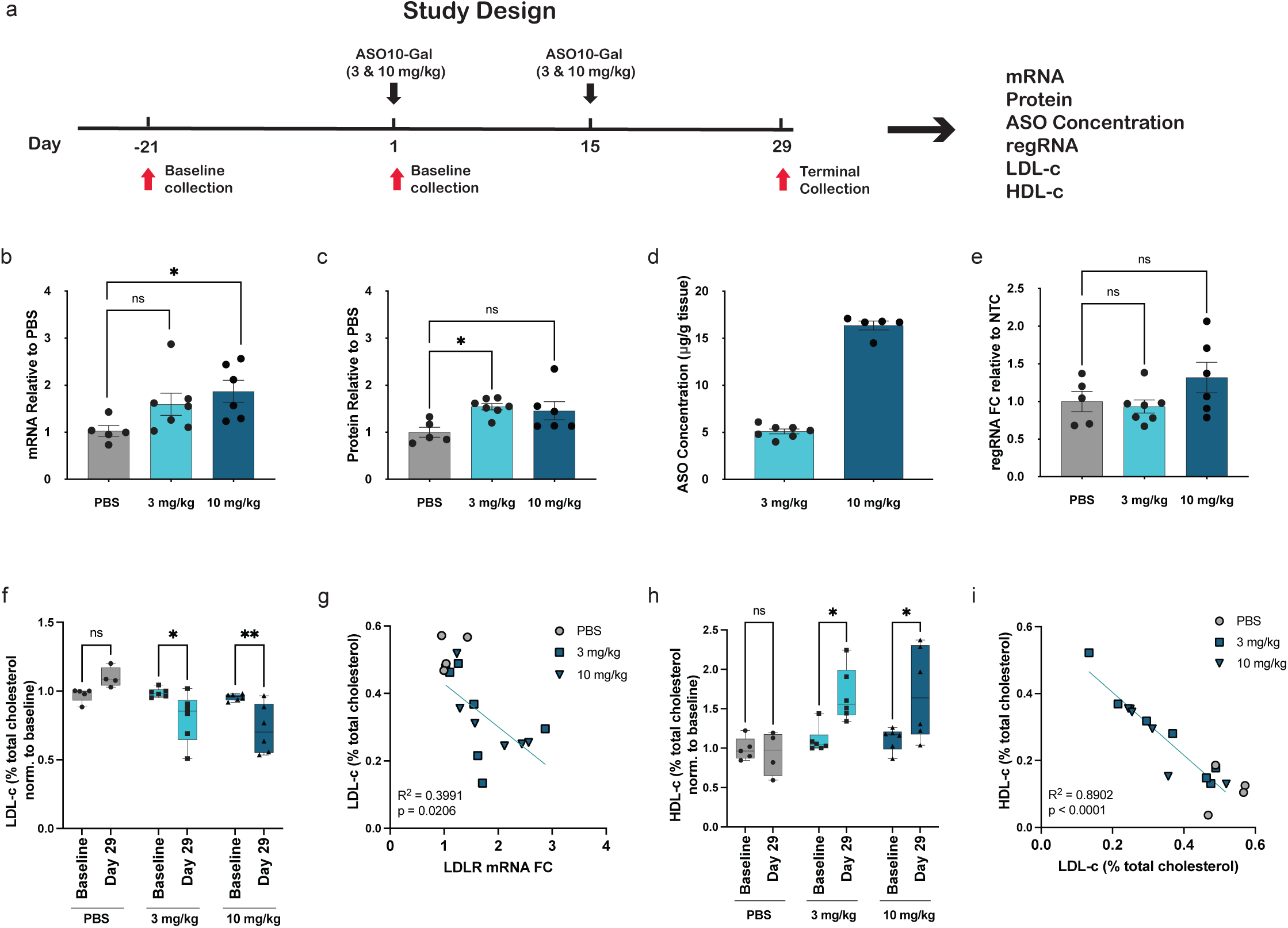
ASO10 upregulates *LDLR* and decreases LDL-c in a humanized liver chimeric mouse model. A) Study design for the PXB^®^ humanized mouse study. Mice (6-month-old male, n= 5-7 per group) received two subcutaneous doses of PBS or GalNAc-conjugated ASO10 at 3 or 10 mg/kg on Day 1 and Day 15 (purple arrows). Blood samples were taken as indicated by red arrows and mice euthanized on Day 29 of study. B) *LDLR* mRNA levels in liver following ASO treatment as assessed by qPCR. Data presented as mean fold change relative to PBS (n= 5-7 per group, one-way ANOVA, * p < 0.05, ns = not significant). C) LDLR protein levels in liver quantified by ELISA. Data presented as mean fold change relative to PBS (n= 5-7 per group, one-way ANOVA, * p < 0.05, ns = not significant). D) ASO10 concentration in liver lysates as determined by hybridization ELISA (n= 5-7 per group). ASO10 was measured but not detected in PBS control livers (not shown). E) *LDLR* regRNA expression in liver as determined by ddPCR. F) Change in LDL-c levels from baseline to Day 29. LDL-c is represented as percentage of total cholesterol (LDL-c/total cholesterol), normalized to individual baseline levels and presented relative to the average of the PBS group (n= 5-7 per group, two-way ANOVA, ** p < 0.01, * p < 0.05, ns = not significant). G) Correlation between *LDLR* mRNA and LDL-c levels. Each point represents an individual mouse (n = 17); R^2^ and p-value determined by simple linear regression and shown on the inset. H) Change in HDL-c levels from baseline to Day 29. HDL-c represented as percentage of total cholesterol (HDL-c/total cholesterol), normalized to individual baseline levels and presented relative to the average of the PBS group (n= 5-7 per group, two-way ANOVA, ** p < 0.01, * p < 0.05, ns = not significant). I) Correlation between LDL and HDL cholesterol levels. Each point represents an individual mouse (n = 17); R^2^ and p-value determined by simple linear regression and shown on the inset.

ASO10-Gal treatment resulted in statistically significant increases in both *LDLR* mRNA and protein levels (Fig. 3B and 3C). A maximal 1.9-fold increase in mRNA was observed at 10 mg/kg, while LDLR protein levels increased 1.5-fold, reaching significance at 3 mg/kg but not at 10 mg/kg (Fig. 3C; note outlier in the 10 mg/kg group). The *LDLR* regRNA levels were quantified, but unlike HepG2 cells, there was not a clear dose-dependent increase beyond a non-significant trend at 10 mg/kg (Fig. 3E). A hybridization-based ELISA confirmed there was a dose-dependent increase in ASO10-Gal concentrations in liver (Fig. 3D).

The functional impact of the observed increase in LDLR was assessed by monitoring circulating LDL-c levels^21^. Relative to PBS control, ASO10-Gal administration resulted in a statistically-significant reduction of LDL-c at both 3 and 10 mg/kg on Day 29 relative to individual Day 1 baseline values (Fig. 3F). At the group level, there was an average 25% decrease in LDL-c with individual mice showing up to a 50% decrease. At the individual animal level, there was a statistically-significant inverse correlation between *LDLR* mRNA and LDL-c levels in response to ASO10-Gal administration, indicating that greater increases in *LDLR* resulted in more substantial reductions in LDL-c (Fig. 3G).

In contrast to LDL-c, which transports cholesterol to tissues, HDL-c carries cholesterol away from tissues and to the liver. LDL-c and HDL-c ratio is often used to assess cardiovascular disease risk with a lower ratio being associated with reduced risk^22^. Both statins and PCSK9 inhibition have been associated with moderately increased HDL-c levels^23,24^. HDL-c levels were therefore evaluated and compared to LDL-c. A statistically-significant 1.5- to 2-fold increase in HDL-c was observed at both dose levels relative to baseline (Fig. 3H), and there was a strong inverse correlation between LDL-c and HDL-c levels for individual animals (Fig. 3I). This result suggests that decreases in LDL-c due to elevated LDLR may be associated with a concomitant shift towards HDL-c that can transport peripheral cholesterol to the liver.

Collectively, these data demonstrate that ASO10-Gal achieves tissue concentrations sufficient to engage the *LDLR* regRNA, increase *LDLR* gene expression, and reduce LDL-c, the major cholesterol species elevated in patients with hypercholesterolemia.

## Discussion

There is a growing recognition that long non-coding RNAs expressed from enhancers and promoters play important roles in transcription regulation^9^. Here, we used epigenomic mapping and regRNA Capture-seq to identify a complex collection of non-coding regRNAs expressed from both the *LDLR* promoter and an active enhancer (Fig. 1). These regRNAs are substantially shorter than the *LDLR* mRNA, largely unspliced, and frequently terminate at canonical polyadenylation signals, consistent with the production of discrete, full-length transcripts rather than degradation intermediates.

Notably, sequences from a subset of the longer regRNAs are conserved between HepG2 cells and human liver tissue, suggesting that their transcription initiation and 3′ end formation are similarly regulated in vivo. In contrast, the shorter RNA species are not conserved between HepG2 cells and liver, suggesting that these isoforms may reflect context-dependent transcription initiation or RNA processing. The conservation of the longer regRNAs from an immortalized cell line to primary human tissue underscores their likely functional importance in transcriptional regulation.

We further demonstrated the functional role of these regRNAs in controlling the expression of *LDLR* both in vitro and in vivo using ASOs targeting regRNA A, one of the more abundant regRNAs identified in HepG2 cells and human liver tissue. RegRNA A is a paRNA expressed in the antisense direction relative to the *LDLR* mRNA and is partially overlapping its TSS. Only a subset of screened ASOs resulted in increased *LDLR* mRNA and protein levels, suggesting that targeting specific regions within the paRNA may be important in regulating *LDLR* transcription. While the precise mechanism of *LDLR* upregulation in response to targeting this paRNA with these ASOs is not yet known, it is clear that cleavage and degradation of the paRNA are not required (Figs. 2E-G). Although initial screens largely identified gapmer ASOs, including ASO10 (Fig. 2B), reduction of the gap length to eliminate the potential for RNase H1 cleavage did not abrogate ASO activity (Figs. 2F-G). Consistent with an RNase H1-independent mechanism of action, ASO10 treatment led to increases in *LDLR* regRNA A levels in vitro (Fig. 2E), further supporting that regRNA cleavage is not required for *LDLR* upregulation. In mice, regRNA A upregulation did not achieve statistical significance (Fig. 3E), potentially reflecting temporal differences between treatment paradigms in cells as compared to in mice, or context-dependent effects on regRNA levels.

It is not yet known how the mechanism of action for ASOs described herein compares to other oligo-mediated upregulation strategies. For example, multi-exonic, several kilobase long natural antisense transcripts (NATs) that traverse entire genes can be targeted with either gapmer or steric ASOs to increase gene expression^24,8,25^. In the case of gapmers, it appears that degradation of an inhibitory NAT is required for upregulation^24^. It is clear, however, that the observed increase in *LDLR* expression does not require the degradation of the *LDLR* paRNA (Fig. 2E-G). Another class of oligonucleotide, small duplex RNAs, or antigene RNAs (agRNAs), have been shown to increase *LDLR* expression through targeting of the *LDLR* paRNA in cells^26^. These agRNAs work in an AGO2-dependent manner in the nucleus but, like ASOs described here, did not decrease paRNA levels^17^. Rather, the agRNAs’ mechanism targets the *LDLR* paRNA without initiating cleavage and recruits transcriptional machinery to increase *LDLR* transcription. It is therefore possible that different classes of noncoding RNAs, as well as different types of oligonucleotides, may collectively utilize distinct mechanisms to achieve gene activation.

It has been shown that TFs can bind eRNAs through positively charged amino acid patches adjacent to their DNA-binding domains, and that eRNA/paRNA levels positively correlate with transcriptional activity^27,28^. Based on these observations, we hypothesize that the *LDLR* paRNA may similarly interact with TFs, and that ASO binding to the paRNA perturbs these interactions, potentially altering the balance of activating and repressive regulatory proteins at the promoter. A related mechanism has been described for the *OTC* eRNA, where ASO treatment perturbed paRNA structure and altered TF recruitment to the targeted enhancer to enhance transcription^15^. Importantly, we have shown that this regulatory mechanism is likely operational in vivo by demonstrating that ASO treatment increased *LDLR* expression in a humanized liver mouse model to reduce LDL-c (Fig. 3). The conservation of paRNA sequence species (Fig. 1A) and the translation of efficacy between in vitro and in vivo systems (Fig. 2 and 3) highlights the highly conserved and potentially critical role of these regRNAs in gene regulation.

Moreover, our in vivo data showing that ASO10 increases *LDLR* expression and reduces circulating LDL-c levels (Fig. 3) supports the concept that targeting the *LDLR* paRNA directly may have potential therapeutic benefit for patients with HeFH. The current standard of care primarily relies on statins and either antibodies or siRNA targeting proprotein convertase subtilsin/kexin type 9 (PCSK9), a protein that when inhibited or knocked down, results in enhanced levels of LDLR at the membrane and an associated increase in LDL-c uptake^29,30^. Multiple other LDL-c lowering therapeutics are in development, including gene editing of *PCSK9* and oral macrocyclic peptides targeting PCSK9^31,32^. Given the many approaches available or in development, it is possible that increasing *LDLR* levels with an *LDLR* regRNA-targeting ASO could find utility in combination with other modalities, or for patients refractory to current treatments.

Taken together, our data support the exploration of ASOs as a potential therapeutic modality to target nuclear noncoding regulatory RNAs to increase expression of protein coding genes for haploinsufficient or loss-of-function diseases. The apparent broad association of regRNA transcription with the transcription of active protein-coding genes suggests that many genes may be amenable to ASO-mediated transcriptional upregulation^13,15,28^. Haploinsufficient diseases, where there is an approximately 50% reduction in healthy protein levels, are particularly well-suited to this approach where modest increases in expression can have a significant impact on disease^33^. As a field, precision nucleic-based therapies may open new avenues for addressing diseases where upregulation could provide patient benefit.

## Methods

### ASO design and synthesis

ASO sequences were designed as 20 nucleotide sequences with a full phosphorothioate backbone, tiled across the full length of *LDLR* regRNA A. Highly repetitive regions and sequences that have less than three nucleotide mismatches to other transcribed regions were excluded. Gapmer and steric versions of each ASO sequence were synthesized using solid-phase synthesis and purified by high-performance liquid chromatography (HPLC). Gapmer ASOs possess the 2′-O-methoxyethyl (2′-MOE) modification on the five terminal nucleotides flanking a central 10-nucleotide DNA core, while steric ASOs have 2’-MOE modifications on each nucleotide. ASO for the in vivo study was synthesized using typical phosphoramidite chemistry in the 3′ to 5′ direction on a custom NittoPhase^™^ solid support (Kinovate Life Sciences, Inc.) loaded with an triantennary N-acetylgalactosamine (GalNAc) ligand, conjugating to the 3′ end of the completed strands.

### Cell culture

HepG2 cells (#HB-8065, American Type Culture Collection) were cultured in DMEM (#11965092, Thermo Fisher Scientific Inc.) supplemented with 10% FBS (#A5256801, Thermo Fisher Scientific Inc.) and 1% Penicillin-Streptomycin (10,000 U/mL) (#15140122, Thermo Fisher Scientific Inc.). Cells were maintained in a humidified incubator at 37 °C with 5% CO2. Reverse transfection was performed at the time of seeding using Lipofectamine^™^ RNAiMAX Transfection Reagent (#13778100, Thermo Fisher Scientific Inc.) diluted in Opti-MEM^™^ I Reduced Serum Medium (#31985062, Thermo Fisher Scientific Inc.) following manufacturer’s recommendations and cells were harvested 48 hours after transfection.

### HepG2 Precision run-on sequencing (PRO-seq)

PRO-seq libraries were generated with HepG2 cells using established protocols^34^ and data were processed as previously described^15^.

### HepG2 Assay for transposase-accessible chromatin using sequencing (ATAC-seq)

HepG2 cells were permeabilized and tagmented using Tn5 transposase following previously established protocols^35^. ATAC-seq libraries were constructed, sequenced and processed as previously described^15^.

### regRNA Capture-seq in HepG2 and human liver

HepG2 RNA was isolated using the RNeasy^®^ Mini Kit (#74104, Qiagen N.V.) following the manufacturer’s instructions. Human liver tissue sample was obtained and RNA was isolated using TRIzol^™^ reagent (#15596026, Thermo Fisher Scientific Inc.) according to the manufacturer’s protocol. Capture probe sets targeting the *LDLR* regRNA were designed and synthesized (myBaits^®^ Custom Hybridization Capture Kit; Daicel Arbor Biosciences) (Supplemental Table 3). RegRNA Capture-seq was performed as previously described^15^.

High-fidelity (HiFi) reads were processed using command-line tools from the SMRT Link suite (v25.3; Pacific Biosciences of California, Inc.) and SQANTI3 (v5.5.1), as previously described^15,36^. Annotated isoforms were displayed in a strand-specific manner for visualizing gene tracks, and a custom R script was used to generate base-level coverage profiles from stacked transcript alignments to visualize signal intensity across each locus.

### RNA quantitation by quantitative PCR (qPCR)

Total RNA was extracted using the RNeasy^®^ 96 Kit (#74182, Qiagen N.V.) following the manufacturer’s instructions and cDNA was generated using the High-Capacity cDNA Reverse Transcription Kit (#4368813, Thermo Fisher Scientific Inc.) following the manufacturer’s instructions. Quantitative PCR was performed using TaqMan^™^ Fast Advanced Master Mix (#4444965, Thermo Fisher Scientific Inc.) on a QuantStudio^™^ 7 Pro Real-Time PCR System (Thermo Fisher Scientific Inc.). Primer sequences for target and control loci are listed in Supplemental Table 4. Enrichment was calculated as percent input or normalized to a reference region using the ΔΔCt method. Data were visualized using Prism software version 10.4.1 (GraphPad Software) with expression values plotted as fold change (FC) mean ± standard deviation (SD) relative to non-targeting control. Relative levels of *LDLR* regRNA in mouse liver were quantified using crystal digital PCR as previously described^15^. Custom expression detection assays were used to quantify regRNA (Supplemental Table 4).

### LDLR protein detection

LDLR protein expression was assessed using the Human LDLR ELISA Kit (#ab270212, Abcam Limited) and analyzed following the manufacturer’s instructions. Data was normalized to total protein as determined using the Pierce^™^ Bicinchoninic Acid (BCA) Protein Assay Kit (#23225, Thermo Fisher Scientific Inc.). LDLR protein obtained from mouse liver samples was assessed using the Human LDLR Quantikine ELISA Kit (#DLDLR0, R&D Systems, Inc.) following the manufacturer’s instructions and normalized to total protein as described above.

### Humanized liver mouse study

Fifteen male 6-month-old humanized liver mice with the cDNA-urokinase-type plasminogen activator/SCID background (PXB-Mouse^®^) were obtained from PhoenixBio Co. Ltd. ^37,38^.

Repopulation of human hepatocytes in the mouse livers was validated by quantifying human serum albumin levels by manufacturer.

Mice were housed at AAALAC-accredited facilities in sterile polyethylene terephthalate (PET) plastic, BPA-free disposable caging (Innovive, LLC) with contact bedding. General procedures for animal care and housing were performed in accordance with the guidelines of study facilities’ Institutional Animal Care and Use Committee (IACUC). Animals were provided with Teklad Global 18% Protein Rodent Diet 2918 (Envigo Bioproducts Inc.) or ProLab^®^ RMH3000 diet (PMI Nutrition International LLC) and laboratory-grade acidified water with or without vitamin C *ad libitum*. Mice were also provided with vitamin C supplements and diet gel as needed.

Mice received two subcutaneous doses of either 3 mg/kg or 10 mg/kg of ASO10-Gal, or phosphate-buffered saline (PBS), on Days 1 and 15 and were euthanized on Day 29. Serum was processed from whole blood collected 21 days prior to the first dose, the day of administration of the first dose (pre-dose/pre-euthanasia) and at the terminal timepoint. Prior to each blood collection including euthanasia, mice were fasted for 4-6 hours.

### Cholesterol profiling

Profiling of total cholesterol and cholesterol within lipoproteins was performed by IDEXX BioAnalytics (North Grafton, MA). Beckman Coulter AU analyzers were used to quantify LDL-cholesterol (#OSR6296, Beckman Coulter, Inc.), HDL-cholesterol (#OSR6295, Beckman Coulter, Inc.), and total cholesterol (#OSR6216, Beckman Coulter, Inc.). Baseline levels of LDL-c and HDL-c were calculated as the average of Day -21 and Day 1 pre-dose levels.

### ASO quantification by hybridization-ELISA

Harvested mouse liver tissue was lysed, and the ASO was quantified using a hybridization-ELISA. The capture probe was a 5’-digoxigenin (Dig)- and 3’-biotin-triethylene glycol (BioTEG)-modified complementary strand immobilized to a neutravidin coted microplate. After hybridization of the ASO and capture probe, unbound capture probe was cleaved using Nuclease P1. Fluorescence of intact capture probe was measured using the AttoPhos^®^ System (Promega Corporation) according to manufacturer’s recommendations at 435/555 nm (excitation/emission). Sample concentrations were determined by interpolation against a standard curve (0.2 to 8 µg/g tissue), which was fit using a four-parameter logistic (4PL) model.

### Statistical analysis

Data are presented as mean ± standard deviation with error bars representing standard deviation. Exact n values correspond to independent biological replicates or individual animals for in vivo studies and are reported explicitly in figure legends; technical replicates were averaged prior to analysis and not treated as independent observations. For qRT-PCR and protein quantification assays (including ELISA and western blot–based measurements), comparisons between two groups were performed using unpaired two-tailed Student’s t-tests, while experiments involving multiple conditions were analyzed using one-way or two-way ANOVA, as appropriate, with post hoc multiple-comparison tests. Animal studies were analyzed using one-way ANOVA for single-factor comparisons and two-way ANOVA for multi-factor analyses. Correlations were assessed using simple linear regression with reporting of R² and associated p-values. GraphPad Prism software version 10.4.1 was used for all analyses with a significance level of 0.05.

### Data availability

Information for publicly available datasets used can be found in Supplementary Table 5.

## Acknowledgements

The authors thank Rachel Meyers and Mary Wertz for helpful comments on the manuscript, and Bryan Matthews and Yuting Liu for technical support. The research was funded by CAMP4 Therapeutics Corporation.

