## Supplemental information for "Antisense Oligonucleotides Targeting an *LDLR* Regulatory RNA Increase *LDLR* Expression and Reduce LDL-cholesterol in Vivo"

**Supplemental Figure Legends**

**Supplemental Fig. 1.** Summary of Capture-seq mRNA reads.

A) Signal tracks at the *LDLR* locus in HepG2 cells and human liver showing a subset of captured mRNA species. ATAC-seq (light green) and PRO-seq (teal) peaks are displayed as reads per million (RPM). PRO-seq (+) and PRO-seq (–) reads are shown above and below the y-axis, respectively. Due to the high number of captured transcripts, only a representative selection of positive-strand Capture-seq mRNA reads (blue) are displayed. Raw read abundance is indicated by the dark-to-light color scale in the legend.

**Supplemental Fig. 2**. Multiple ASOs upregulate *LDLR* gene expression.

A) *LDLR* fold change of 44 sequences screened as both gapmer ASOs (ASO1-44) and B) steric ASOs (ASO45-88) targeting the *LDLR* regRNA. HepG2 cells were transfected with 100 nM ASO for 48 hours and assessed for *LDLR* expression using qRT-PCR. Data are presented as mean fold change relative to respective NTC ± s.d. (n = 3). ASO hits are highlighted in dark blue.

C) Validation of all primary screen hits. HepG2 cells were transfected with 11nM , 33 nM or 100 nM of ASO for 48 hours. *LDLR* expression was assessed using qRT-PCR (n=3). Data are presented as mean fold change relative to respective NTC ± s.d. (n = 3).

Supp Fig 1

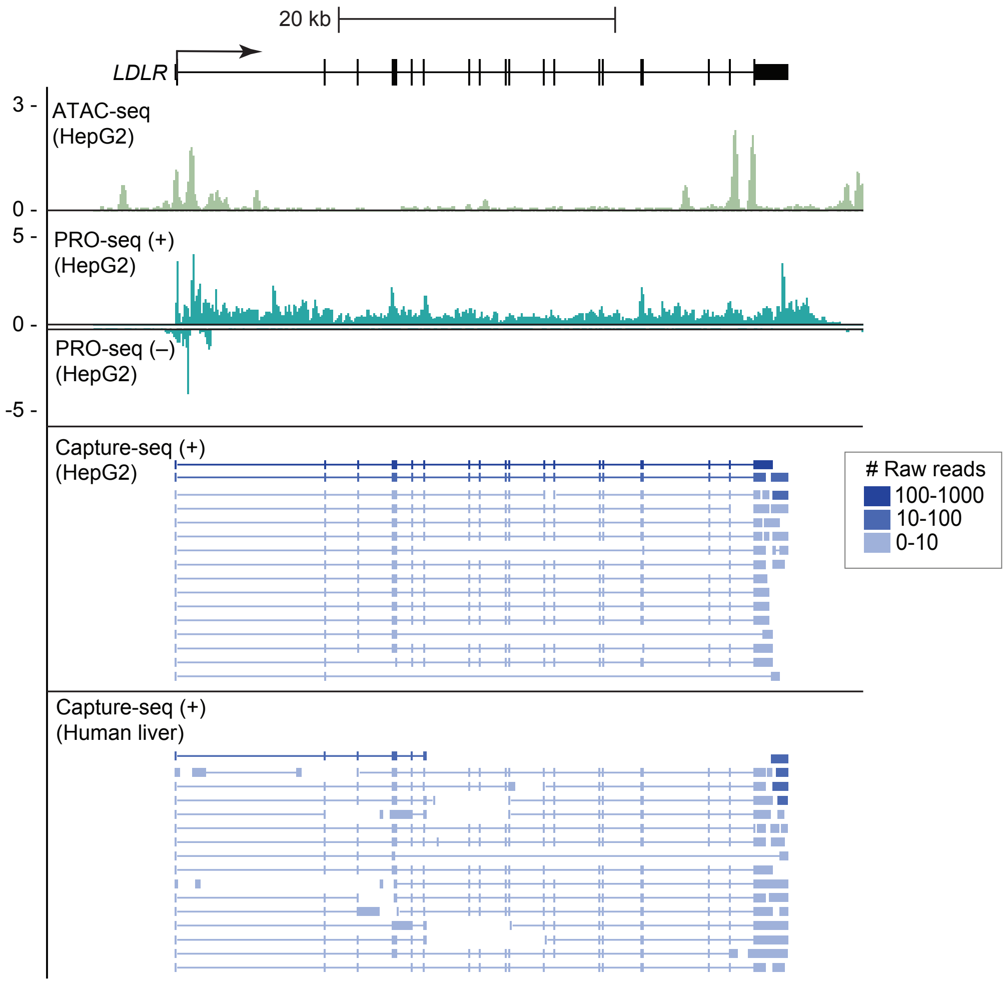

20 kb
LDLR
3 -
ATAC-seq
(HepG2)
0 -
5 -
PRO-seq (+)
(HepG2)
0 -
PRO-seq (–)
(HepG2)
-5 -
Capture-seq (+)
(HepG2)
### Raw reads
100-1000
10-100
0-10
Capture-seq (+)
(Human liver)

Supp Fig 2

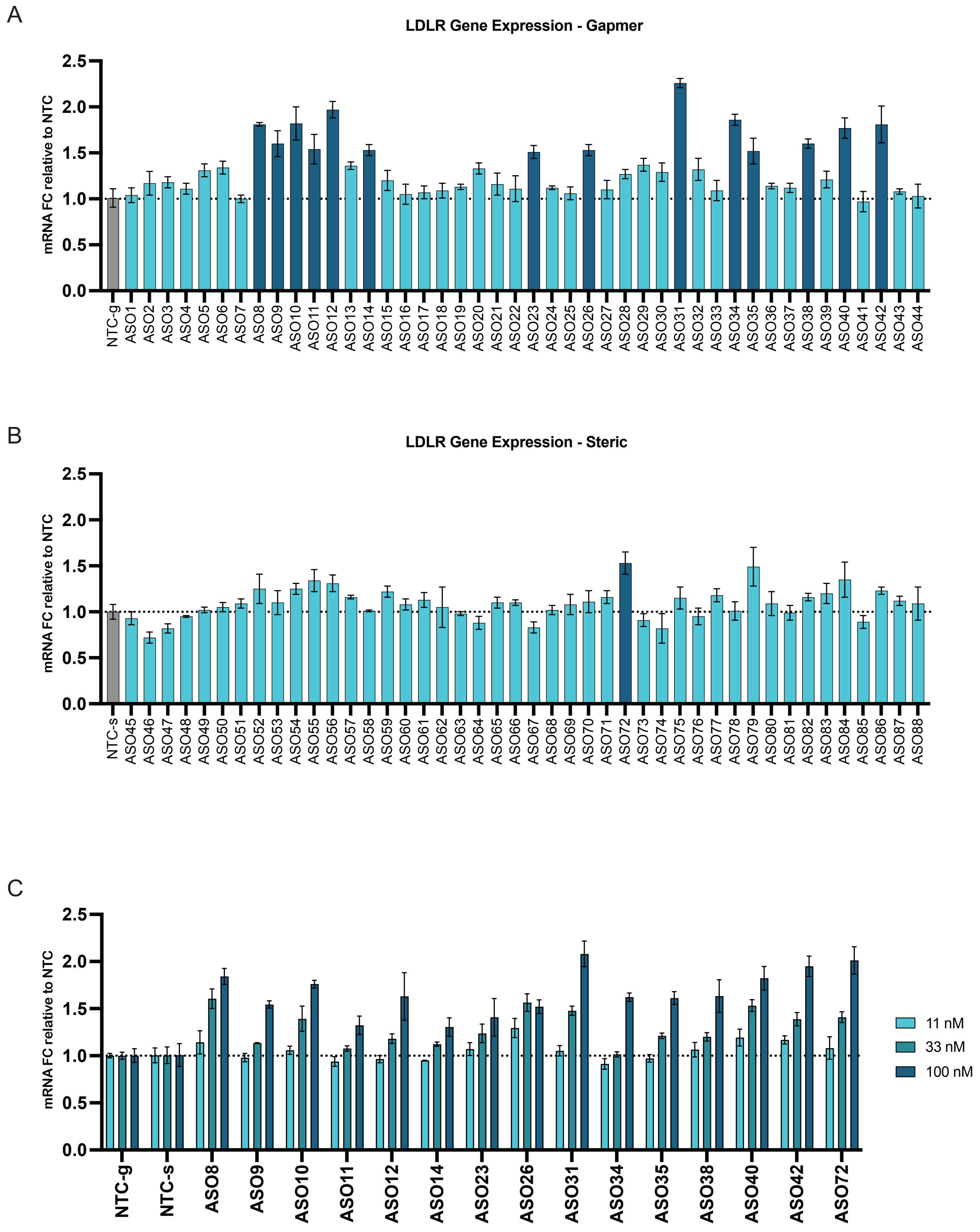
A
LDLR Gene Expression - Gapmer
mRNA FC relative to NTC
2.5
2.0
1.5
1.0
0.5
0.0
NTC-g
ASO1
ASO2
ASO3
ASO4
ASO5
ASO6
ASO7
ASO8
ASO9
ASO10
ASO11
ASO12
ASO13
ASO14
ASO15
ASO16
ASO17
ASO18
ASO19
ASO20
ASO21
ASO22
ASO23
ASO24
ASO25
ASO26
ASO27
ASO28
ASO29
ASO30
ASO31
ASO32
ASO33
ASO34
ASO35
ASO36
ASO37
ASO38
ASO39
ASO40
ASO41
ASO42
ASO43
ASO44
B
LDLR Gene Expression - Steric
mRNA FC relative to NTC
2.5
2.0
1.5
1.0
0.5
0.0
NTC-s
ASO45
ASO46
ASO47
ASO48
ASO49
ASO50
ASO51
ASO52
ASO53
ASO54
ASO55
ASO56
ASO57
ASO58
ASO59
ASO60
ASO61
ASO62
ASO63
ASO64
ASO65
ASO66
ASO67
ASO68
ASO69
ASO70
ASO71
ASO72
ASO73
ASO74
ASO75
ASO76
ASO77
ASO78
ASO79
ASO80
ASO81
ASO82
ASO83
ASO84
ASO85
ASO86
ASO87
ASO88
C
mRNA FC relative to NTC
2.5
2.0
1.5
1.0
0.5
0.0
NTC-g
NTC-s
ASO8
ASO9
ASO10
ASO11
ASO12
ASO14
ASO23
ASO26
ASO31
ASO34
ASO35
ASO38
ASO40
ASO42
ASO72
11 nM
33 nM
100 nM

#### Supplemental Tables

**Supplemental Table 1: Characterized regRNA coordinates.**

| **Annotated *LDLR* regRNA** | **Genome** | **Coordinates** |
|---|---|---|
| Matsui et al. | hg38 | chr19:11088922-11090283 |
| Ray et al. | hg38 | chr19:11089069-11089659 |
| regRNA A | hg38 | chr19:11088975-11090364 |
| regRNA B | hg38 | chr19:11088890-11091963 |
| regRNA C | hg38 | chr19:11089648-11090378 |
| regRNA D | hg38 | chr19:11091407-11091979 |

**Supplemental Table 2: Screened ASO sequences.**

| **Gapmer ASO #** | **Steric ASO #** | **Sequence** |
|---|---|---|
| ASO1 | ASO45 | GCCACTGCCATCCTGAGTTG |
| ASO2 | ASO46 | CTAACTCGGAGCTGGGTGTG |
| ASO3 | ASO47 | CTTCTAACTCGGAGCTGGGT |
| ASO4 | ASO48 | ACCCTCTCTTCTAACTCGGA |
| ASO5 | ASO49 | GGGACCCTCTCTTCTAACTC |
| ASO6 | ASO50 | CAGGTGATCTGCCCAAAAGT |
| ASO7 | ASO51 | GGGACCGCATCTTCTTTGAG |
| ASO8 | ASO52 | GAGGGATTTAAGGGAAACGG |
| ASO9 | ASO53 | CTGAGGGATTTAAGGGAAAC |
| ASO10 | ASO54 | GTCTGAGGGATTTAAGGGAA |
| ASO11 | ASO55 | GGGAGGAGTCTGAGGGATTT |
| ASO12 | ASO56 | AGAATAGGTTGAGAGGGAGC |
| ASO13 | ASO57 | GGAAATTGCGCTGGACCGTC |
| ASO14 | ASO58 | ACAGCAGGTCGTGATCCGGG |
| ASO15 | ASO59 | CAGCTAGGACACAGCAGGTC |
| ASO16 | ASO60 | GAGTGCAATCGCGGGAAGCC |
| ASO17 | ASO61 | CCCTGCTAGAAACCTCACAT |
| ASO18 | ASO62 | TCCCCCTGCTAGAAACCTCA |
| ASO19 | ASO63 | GGATCTTTCAGAAGATGCGT |
| ASO20 | ASO64 | AGGCTATTGGAGGATCTTGA |
| ASO21 | ASO65 | GAGGACAATGGCATTAGGCT |
| ASO22 | ASO66 | AGTGGCGGAAGTTCCCAACA |

| | | |
|---|---|---|
| ASO23 | ASO67 | TGGAGACCCGGATGTCCAGC |
| ASO24 | ASO68 | ACCCCCGCCTTAAGTCCTTC |
| ASO25 | ASO69 | CTAAAACAAGGGATCACTCC |
| ASO26 | ASO70 | GATCTGCCCAAAAGTGCTGG |
| ASO27 | ASO71 | GTCCCCCAAGTCTCCACAGG |
| ASO28 | ASO72 | GGTCCTTCCCAAACGGCTGG |
| ASO29 | ASO73 | CATCCCTGGGAGACTTGTGG |
| ASO30 | ASO74 | GGCGCCAGAATAGGTTGAGA |
| ASO31 | ASO75 | GGGACTGCAGGTAAGGCTTG |
| ASO32 | ASO76 | TGGACCGTCGCCTTGCTCCT |
| ASO33 | ASO77 | AGGACACAGCAGGTCGTGAT |
| ASO34 | ASO78 | CAATCGCGGGAAGCCAGGGT |
| ASO35 | ASO79 | AATGCTGTAAATGACGTGGG |
| ASO36 | ASO80 | ACTCCTCCCCCTGCTAGAAA |
| ASO37 | ASO81 | GGCCGTTCGAAACTCCTCCT |
| ASO38 | ASO82 | CGAAGGACTGGAGTGGGAAT |
| ASO39 | ASO83 | GCGTCAGCTCTTCACCGGAG |
| ASO40 | ASO84 | TGCGTTTCCAATTTTGAGGG |
| ASO41 | ASO85 | CGAGAAATTTCAGGAGGATC |
| ASO42 | ASO86 | GCATTAGGCTATTGGAGGAT |
| ASO43 | ASO87 | TTTGAGGCAGAGAGGACAAT |
| ASO44 | ASO88 | TAGAAAGTGGCGGAAGTTCC |
| NTC-g | NTC-s | AAGGTTCGGAGTCGTAGCTG |

**Supplemental Table 3: Capture-seq Probe Sequences.**

| **Probe ID** | **Probe Sequence (5'-3')** |
|---|---|
| chr19:11088475-11090864_0 | AGAGGTTAGCACTGCCTGCCAAAGTCAGTTTGCAAAATCCCAGAGAAATCCAGCTTATTCCTGGGGGAACCGCCAAGACT |
| chr19:11088475-11090864_21 | AAGTCAGTTTGCAAAATCCCAGAGAAATCCAGCTTATTCCTGGGGGAACCGCCAAGACTGCCCAGCCCTGTGTGGGGTTC |
| chr19:11088475-11090864_42 | GAGAAATCCAGCTTATTCCTGGGGGAACCGCCAAGACTGCCCAGCCCTGTGTGGGGTTCAGGCAAGTTTCTCACATGTGC |
| chr19:11088475-11090864_63 | GGGGAACCGCCAAGACTGCCCAGCCCTGTGTGGGGTTCAGGCAAGTTTCTCACATGTGCCTTTTTGGCAAGAGGCCTCTG |
| chr19:11088475-11090864_84 | AGCCCTGTGTGGGGTTCAGGCAAGTTTCTCACATGTGCCTTTTTGGCAAGAGGCCTCTGGCAACCCCATGAGTCCCCAAA |
| chr19:11088475-11090864_105 | AAGTTTCTCACATGTGCCTTTTTGGCAAGAGGCCTCTGGCAACCCCATGAGTCCCCAAAGAGACTCAATTCTAAAAGTTG |
| chr19:11088475-11090864_126 | TTGGCAAGAGGCCTCTGGCAACCCCATGAGTCCCCAAAGAGACTCAATTCTAAAAGTTGGTCTCCACCAGCTCTCTGTGG |
| chr19:11088475-11090864_147 | CCCCATGAGTCCCCAAAGAGACTCAATTCTAAAAGTTGGTCTCCACCAGCTCTCTGTGGCTTAGGGGTTCAAGTTCAACT |

| | |
|---|---|
| chr19:11088475-11090864_168 | CTCAATTCTAAAAGTTGGTCTCCACCAGCTCTCTGTGGCTTAGGGGTTCAAGTTCAACTGTGAAAGCCCTGTTTTGTTTT |
| chr19:11088475-11090864_189 | CCACCAGCTCTCTGTGGCTTAGGGGTTCAAGTTCAACTGTGAAAGCCCTGTTTTGTTTTGATTTTGCTTTGAGGGAGAGG |
| chr19:11088475-11090864_210 | GGGGTTCAAGTTCAACTGTGAAAGCCCTGTTTTGTTTTGATTTTGCTTTGAGGGAGAGGAAACCGCCCTTCTGTTTGTTC |
| chr19:11088475-11090864_231 | AAGCCCTGTTTTGTTTTGATTTTGCTTTGAGGGAGAGGAAACCGCCCTTCTGTTTGTTCAACTCCTTCTCCTAAGGGGAG |
| chr19:11088475-11090864_252 | TTGCTTTGAGGGAGAGGAAACCGCCCTTCTGTTTGTTCAACTCCTTCTCCTAAGGGGAGAAATCAATATTTACGTCCAGA |
| chr19:11088475-11090864_273 | CGCCCTTCTGTTTGTTCAACTCCTTCTCCTAAGGGGAGAAATCAATATTTACGTCCAGACTCCAGGTATCCGTACAATTG |
| chr19:11088475-11090864_294 | CCTTCTCCTAAGGGGAGAAATCAATATTTACGTCCAGACTCCAGGTATCCGTACAATTGATTTTTCAGATGTTTATACTC |
| chr19:11088475-11090864_315 | CAATATTTACGTCCAGACTCCAGGTATCCGTACAATTGATTTTTCAGATGTTTATACTCAGCCAAAGGCGGGATCCCACA |
| chr19:11088475-11090864_336 | AGGTATCCGTACAATTGATTTTTCAGATGTTTATACTCAGCCAAAGGCGGGATCCCACAAAACAAAAAATATTTTTTTGG |
| chr19:11088475-11090864_357 | TTCAGATGTTTATACTCAGCCAAAGGCGGGATCCCACAAAACAAAAAATATTTTTTTGGCTGTACTTTTGTGAAGATTTT |
| chr19:11088475-11090864_378 | AAAGGCGGGATCCCACAAAACAAAAAATATTTTTTTGGCTGTACTTTTGTGAAGATTTTATTTAAATTCCTGATTGATCA |
| chr19:11088475-11090864_399 | AAAAAATATTTTTTTGGCTGTACTTTTGTGAAGATTTTATTTAAATTCCTGATTGATCAGTGTCTATTAGGTGATTTGGA |
| chr19:11088475-11090864_420 | ACTTTTGTGAAGATTTTATTTAAATTCCTGATTGATCAGTGTCTATTAGGTGATTTGGAATAACAATGTAAAAACAATAT |
| chr19:11088475-11090864_441 | AAATTCCTGATTGATCAGTGTCTATTAGGTGATTTGGAATAACAATGTAAAAACAATATACAACGAAAGGAAGCTAAAAA |
| chr19:11088475-11090864_462 | CTATTAGGTGATTTGGAATAACAATGTAAAAACAATATACAACGAAAGGAAGCTAAAAATCTATACACAATTCCTAGAAA |
| chr19:11088475-11090864_483 | CAATGTAAAAACAATATACAACGAAAGGAAGCTAAAAATCTATACACAATTCCTAGAAAGGAAAAGGCAAATATAGAAAG |
| chr19:11088475-11090864_504 | CGAAAGGAAGCTAAAAATCTATACACAATTCCTAGAAAGGAAAAGGCAAATATAGAAAGTGGCGGAAGTTCCCAACATTT |
| chr19:11088475-11090864_525 | TACACAATTCCTAGAAAGGAAAAGGCAAATATAGAAAGTGGCGGAAGTTCCCAACATTTTTAGTGTTTTCCTTTTGAGGC |
| chr19:11088475-11090864_546 | AAGGCAAATATAGAAAGTGGCGGAAGTTCCCAACATTTTTAGTGTTTTCCTTTTGAGGCAGAGAGGACAATGGCATTAGG |
| chr19:11088475-11090864_567 | GGAAGTTCCCAACATTTTTAGTGTTTTCCTTTTGAGGCAGAGAGGACAATGGCATTAGGCTATTGGAGGATCTTGAAAGG |
| chr19:11088475-11090864_588 | TGTTTTCCTTTTGAGGCAGAGAGGACAATGGCATTAGGCTATTGGAGGATCTTGAAAGGCTGTTGTTATCCTTCTGTGGA |
| chr19:11088475-11090864_609 | AGGACAATGGCATTAGGCTATTGGAGGATCTTGAAAGGCTGTTGTTATCCTTCTGTGGACAACAACAGCAAAATGTTAAC |
| chr19:11088475-11090864_630 | TGGAGGATCTTGAAAGGCTGTTGTTATCCTTCTGTGGACAACAACAGCAAAATGTTAACAGTTAAACATCGAGAAATTTC |

| | |
|---|---|
| chr19:11088475-11090864_651 | TGTTATCCTTCTGTGGACAACAACAGCAAAATGTTAACAGTTAAACATCGAGAAATTTCAGGAGGATCTTTCAGAAGATG |
| chr19:11088475-11090864_672 | AACAGCAAAATGTTAACAGTTAAACATCGAGAAATTTCAGGAGGATCTTTCAGAAGATGCGTTTCCAATTTTGAGGGGGC |
| chr19:11088475-11090864_693 | AAACATCGAGAAATTTCAGGAGGATCTTTCAGAAGATGCGTTTCCAATTTTGAGGGGGCGTCAGCTCTTCACCGGAGACC |
| chr19:11088475-11090864_714 | GGATCTTTCAGAAGATGCGTTTCCAATTTTGAGGGGGCGTCAGCTCTTCACCGGAGACCCAAATACAACAAATCAAGTCG |
| chr19:11088475-11090864_735 | TCCAATTTTGAGGGGGCGTCAGCTCTTCACCGGAGACCCAAATACAACAAATCAAGTCGCCTGCCCTGGCGACACTTTCG |
| chr19:11088475-11090864_756 | GCTCTTCACCGGAGACCCAAATACAACAAATCAAGTCGCCTGCCCTGGCGACACTTTCGAAGGACTGGAGTGGGAATCAG |
| chr19:11088475-11090864_777 | TACAACAAATCAAGTCGCCTGCCCTGGCGACACTTTCGAAGGACTGGAGTGGGAATCAGAGCTTCACGGGTTAAAAAGCC |
| chr19:11088475-11090864_798 | CCCTGGCGACACTTTCGAAGGACTGGAGTGGGAATCAGAGCTTCACGGGTTAAAAAGCCGATGTCACATCGGCCGTTCGA |
| chr19:11088475-11090864_819 | ACTGGAGTGGGAATCAGAGCTTCACGGGTTAAAAAGCCGATGTCACATCGGCCGTTCGAAACTCCTCCTCTTGCAGTGAG |
| chr19:11088475-11090864_840 | TCACGGGTTAAAAAGCCGATGTCACATCGGCCGTTCGAAACTCCTCCTCTTGCAGTGAGGTGAAGACATTTGAAAATCAC |
| chr19:11088475-11090864_861 | TCACATCGGCCGTTCGAAACTCCTCCTCTTGCAGTGAGGTGAAGACATTTGAAAATCACCCCACTGCAAACTCCTCCCCC |
| chr19:11088475-11090864_882 | CCTCCTCTTGCAGTGAGGTGAAGACATTTGAAAATCACCCCACTGCAAACTCCTCCCCCTGCTAGAAACCTCACATTGAA |
| chr19:11088475-11090864_903 | AGACATTTGAAAATCACCCCACTGCAAACTCCTCCCCCTGCTAGAAACCTCACATTGAAATGCTGTAAATGACGTGGGCC |
| chr19:11088475-11090864_924 | CTGCAAACTCCTCCCCCTGCTAGAAACCTCACATTGAAATGCTGTAAATGACGTGGGCCCCGAGTGCAATCGCGGGAAGC |
| chr19:11088475-11090864_945 | AGAAACCTCACATTGAAATGCTGTAAATGACGTGGGCCCCGAGTGCAATCGCGGGAAGCCAGGGTTTCCAGCTAGGACAC |
| chr19:11088475-11090864_966 | TGTAAATGACGTGGGCCCCGAGTGCAATCGCGGGAAGCCAGGGTTTCCAGCTAGGACACAGCAGGTCGTGATCCGGGTCG |
| chr19:11088475-11090864_987 | GTGCAATCGCGGGAAGCCAGGGTTTCCAGCTAGGACACAGCAGGTCGTGATCCGGGTCGGGACACTGCCTGGCAGAGGCT |
| chr19:11088475-11090864_1008 | GTTTCCAGCTAGGACACAGCAGGTCGTGATCCGGGTCGGGACACTGCCTGGCAGAGGCTGCGAGCATGGGGCCCTGGGGC |
| chr19:11088475-11090864_1029 | GGTCGTGATCCGGGTCGGGACACTGCCTGGCAGAGGCTGCGAGCATGGGGCCCTGGGGCTGGAAATTGCGCTGGACCGTC |
| chr19:11088475-11090864_1050 | ACTGCCTGGCAGAGGCTGCGAGCATGGGGCCCTGGGGCTGGAAATTGCGCTGGACCGTCGCCTTGCTCCTCGCCGCGGCG |
| chr19:11088475-11090864_1071 | GCATGGGGCCCTGGGGCTGGAAATTGCGCTGGACCGTCGCCTTGCTCCTCGCCGCGGCGGGGACTGCAGGTAAGGCTTGC |
| chr19:11088475-11090864_1092 | AATTGCGCTGGACCGTCGCCTTGCTCCTCGCCGCGGCGGGGACTGCAGGTAAGGCTTGCTCCAGGCGCCAGAATAGGTTG |
| chr19:11088475-11090864_1113 | TGCTCCTCGCCGCGGCGGGGACTGCAGGTAAGGCTTGCTCCAGGCGCCAGAATAGGTTGAGAGGGAGCCCCCGGGGGGCC |

| | |
|---|---|
| chr19:11088475-11090864_1134 | CTGCAGGTAAGGCTTGCTCCAGGCGCCAGAATAGGTTGAGAGGGAGCCCCCGGGGGGCCCTTGGGAATTTATTTTTTTGG |
| chr19:11088475-11090864_1155 | GGCGCCAGAATAGGTTGAGAGGGAGCCCCCGGGGGGCCCTTGGGAATTTATTTTTTTGGGTACAAATAATCACTCCATCC |
| chr19:11088475-11090864_1176 | GGAGCCCCCGGGGGGCCCTTGGGAATTTATTTTTTTGGGTACAAATAATCACTCCATCCCTGGGAGACTTGTGGGGTAAT |
| chr19:11088475-11090864_1197 | GGAATTTATTTTTTTGGGTACAAATAATCACTCCATCCCTGGGAGACTTGTGGGGTAATGGCACGGGGTCCTTCCCAAAC |
| chr19:11088475-11090864_1218 | AAATAATCACTCCATCCCTGGGAGACTTGTGGGGTAATGGCACGGGGTCCTTCCCAAACGGCTGGAGGGGGCGCTGGAGG |
| chr19:11088475-11090864_1302 | CGCTGAGGGGAGCGCGAGGGTCGGGAGGAGTCTGAGGGATTTAAGGGAAACGGGGCACCGCTGTCCCCCAAGTCTCCACA |
| chr19:11088475-11090864_1323 | CGGGAGGAGTCTGAGGGATTTAAGGGAAACGGGGCACCGCTGTCCCCCAAGTCTCCACAGGGTGAGGGACCGCATCTTCT |
| chr19:11088475-11090864_1659 | GCCACCGCGCCCGGCCGGGACCCTCTCTTCTAACTCGGAGCTGGGTGTGGGGACCTCCAGTCCTAAAACAAGGGATCACT |
| chr19:11088475-11090864_1680 | CCTCTCTTCTAACTCGGAGCTGGGTGTGGGGACCTCCAGTCCTAAAACAAGGGATCACTCCCACCCCCGCCTTAAGTCCT |
| >chr19:11088475-11090864_1701 | GGGTGTGGGGACCTCCAGTCCTAAAACAAGGGATCACTCCCACCCCCGCCTTAAGTCCTTCTGGGGGCGAGGGCGACTGG |
| chr19:11088475-11090864_1722 | TAAAACAAGGGATCACTCCCACCCCCGCCTTAAGTCCTTCTGGGGGCGAGGGCGACTGGAGACCCGGATGTCCAGCCTGG |
| chr19:11088475-11090864_1743 | CCCCCGCCTTAAGTCCTTCTGGGGGCGAGGGCGACTGGAGACCCGGATGTCCAGCCTGGAGGTCACCGCGGGCTCAGGGG |
| chr19:11088475-11090864_1764 | GGGGCGAGGGCGACTGGAGACCCGGATGTCCAGCCTGGAGGTCACCGCGGGCTCAGGGGTCCCGATCCGCTTTGCGCGAC |
| chr19:11088475-11090864_1785 | CCGGATGTCCAGCCTGGAGGTCACCGCGGGCTCAGGGGTCCCGATCCGCTTTGCGCGACCCCAGGGCGCCACTGCCATCC |
| chr19:11088475-11090864_1806 | CACCGCGGGCTCAGGGGTCCCGATCCGCTTTGCGCGACCCCAGGGCGCCACTGCCATCCTGAGTTGGGTGCAGTCCCGGG |
| chr19:11088475-11090864_1827 | GATCCGCTTTGCGCGACCCCAGGGCGCCACTGCCATCCTGAGTTGGGTGCAGTCCCGGGATTCCGCCGCGTGCTCCGGGA |
| chr19:11088475-11090864_1848 | GGGCGCCACTGCCATCCTGAGTTGGGTGCAGTCCCGGGATTCCGCCGCGTGCTCCGGGACGGGGGCCACCCCCTCCCGCC |
| chr19:11088475-11090864_1953 | CCCCGAATTCCATTGGGTGTAGTCCAACAGGCCACCCTCGAGCCACTCCCCTTGTCCAATGTGAGGCGGTGGAGGCGGAG |
| chr19:11088475-11090864_1974 | GTCCAACAGGCCACCCTCGAGCCACTCCCCTTGTCCAATGTGAGGCGGTGGAGGCGGAGGCGGGCGTCGGGAGGACGGGG |
| chr19:11088475-11090864_1995 | CCACTCCCCTTGTCCAATGTGAGGCGGTGGAGGCGGAGGCGGGCGTCGGGAGGACGGGGCTTGTGTACGAGCGGGGCGGG |
| chr19:11088475-11090864_2016 | AGGCGGTGGAGGCGGAGGCGGGCGTCGGGAGGACGGGGCTTGTGTACGAGCGGGGCGGGGCTGGCGCGGAAGTCTGAGCC |
| chr19:11088475-11090864_2037 | GCGTCGGGAGGACGGGGCTTGTGTACGAGCGGGGCGGGGCTGGCGCGGAAGTCTGAGCCTCACCTTGTCCGGGGCGAGGC |
| chr19:11088475-11090864_2058 | TGTACGAGCGGGGCGGGGCTGGCGCGGAAGTCTGAGCCTCACCTTGTCCGGGGCGAGGCGGATGCAGGGGAGGCCTGGCG |

| | |
|---|---|
| chr19:11088475-11090864_2079 | GCGCGGAAGTCTGAGCCTCACCTTGTCCGGGGCGAGGCGGATGCAGGGGAGGCCTGGCGTTCCTCCGCGGTTCCTGTCAC |
| chr19:11088475-11090864_2100 | CTTGTCCGGGGCGAGGCGGATGCAGGGGAGGCCTGGCGTTCCTCCGCGGTTCCTGTCACAAAGGCGACGACAAGTCCCGG |
| chr19:11088475-11090864_2121 | GCAGGGGAGGCCTGGCGTTCCTCCGCGGTTCCTGTCACAAAGGCGACGACAAGTCCCGGGTCCCCGGAGCCGCCTCCGCG |
| chr19:11088475-11090864_2142 | TCCGCGGTTCCTGTCACAAAGGCGACGACAAGTCCCGGGTCCCCGGAGCCGCCTCCGCGACATACACGAGTCGCCCTCCG |
| chr19:11088475-11090864_2163 | GCGACGACAAGTCCCGGGTCCCCGGAGCCGCCTCCGCGACATACACGAGTCGCCCTCCGTTATCCTGGGCCCTCCTGGCG |
| chr19:11088475-11090864_2184 | CCGGAGCCGCCTCCGCGACATACACGAGTCGCCCTCCGTTATCCTGGGCCCTCCTGGCGAAGTCCCCGGTTTCCGCTGTG |
| chr19:11088475-11090864_2205 | ACACGAGTCGCCCTCCGTTATCCTGGGCCCTCCTGGCGAAGTCCCCGGTTTCCGCTGTGCTCTGTGGCGACACCTCCGTC |
| chr19:11088475-11090864_2226 | CCTGGGCCCTCCTGGCGAAGTCCCCGGTTTCCGCTGTGCTCTGTGGCGACACCTCCGTCCCCACCTTGTCCTGGGGGGCG |
| chr19:11088475-11090864_2247 | CCCCGGTTTCCGCTGTGCTCTGTGGCGACACCTCCGTCCCCACCTTGTCCTGGGGGGCGCCCTCGCCCCACCAGCCCCGA |
| chr19:11088475-11090864_2268 | GTGGCGACACCTCCGTCCCCACCTTGTCCTGGGGGGCGCCCTCGCCCCACCAGCCCCGATCAAGTTCACAGAGGGGCCCC |
| chr19:11088475-11090864_2289 | CCTTGTCCTGGGGGGCGCCCTCGCCCCACCAGCCCCGATCAAGTTCACAGAGGGGCCCCCGGCCACCCTCAAGGCCTCGG |
| chr19:11088475-11090864_2309 | TCGCCCCACCAGCCCCGATCAAGTTCACAGAGGGGCCCCCGGCCACCCTCAAGGCCTCGGTTCCTTACGAGGTTGAAACG |
| chr19:11090988-11091987_0 | CGGTTCAGCTGTCCAAGCGGCGATTTTTCCTCTGGGTGAAATGGATTAGATTTTAGATTTCCACAAGAGGCTGGTTAGTG |
| chr19:11090988-11091987_21 | GATTTTTCCTCTGGGTGAAATGGATTAGATTTTAGATTTCCACAAGAGGCTGGTTAGTGCATGATCCTGAGTTAGAGCTT |
| chr19:11090988-11091987_42 | GGATTAGATTTTAGATTTCCACAAGAGGCTGGTTAGTGCATGATCCTGAGTTAGAGCTTTTTAGGTGGCTTTAAATTAGT |
| chr19:11090988-11091987_63 | CAAGAGGCTGGTTAGTGCATGATCCTGAGTTAGAGCTTTTTAGGTGGCTTTAAATTAGTTGCAGAGAGACAGCCTCGCCC |
| chr19:11090988-11091987_84 | ATCCTGAGTTAGAGCTTTTTAGGTGGCTTTAAATTAGTTGCAGAGAGACAGCCTCGCCCTAGACAACAGCTACATGGCCC |
| chr19:11090988-11091987_105 | GGTGGCTTTAAATTAGTTGCAGAGAGACAGCCTCGCCCTAGACAACAGCTACATGGCCCTTTCCCTCCTGAGAACCAGCC |
| chr19:11090988-11091987_126 | GAGAGACAGCCTCGCCCTAGACAACAGCTACATGGCCCTTTCCCTCCTGAGAACCAGCCTAGCCTAGAAAAGGATTGGGA |
| chr19:11090988-11091987_147 | CAACAGCTACATGGCCCTTTCCCTCCTGAGAACCAGCCTAGCCTAGAAAAGGATTGGGATTGCCTGATGAACACAAGGAT |
| chr19:11090988-11091987_168 | CCTCCTGAGAACCAGCCTAGCCTAGAAAAGGATTGGGATTGCCTGATGAACACAAGGATTGCAGGAAACTTTTTTTTTAA |
| chr19:11090988-11091987_189 | CTAGAAAAGGATTGGGATTGCCTGATGAACACAAGGATTGCAGGAAACTTTTTTTTTAATTGGCAAGGGGGTTGGCTTTG |
| chr19:11090988-11091987_210 | CTGATGAACACAAGGATTGCAGGAAACTTTTTTTTTAATTGGCAAGGGGGTTGGCTTTGACTGGATGGAGAGCTTTGAAC |

| | |
|---|---|
| chr19:11090988-11091987_231 | GGAAACTTTTTTTTAATTGGCAAGGGGGTTGGCTTTGACTGGATGGAGAGCTTTGAACTGCCTTGAAATTCACGCTGTA |
| chr19:11090988-11091987_252 | CAAGGGGGTTGGCTTTGACTGGATGGAGAGCTTTGAACTGCCTTGAAATTCACGCTGTAACTAACACACCAGTTTCCTCT |
| chr19:11090988-11091987_273 | GATGGAGAGCTTTGAACTGCCTTGAAATTCACGCTGTAACTAACACACCAGTTTCCTCTGGGAGGCCAGAGAGGGAGGGA |
| chr19:11090988-11091987_294 | TTGAAATTCACGCTGTAACTAACACACCAGTTTCCTCTGGGAGGCCAGAGAGGGAGGGAGGGTGTAATGAAATACGGATG |
| chr19:11090988-11091987_315 | ACACACCAGTTTCCTCTGGGAGGCCAGAGAGGGAGGGAGGGTGTAATGAAATACGGATGATTGTTCTTTTATTTTTATTT |
| chr19:11090988-11091987_525 | CCGCGCCCCACCGGGGATGATGATGATTGCAAACATTCTGCCACTCAGTTTTACAAAAGAAAGAGAGGCACTGGATTAAT |
| chr19:11090988-11091987_546 | GATGATTGCAAACATTCTGCCACTCAGTTTTACAAAAGAAAGAGAGGCACTGGATTAATGTGTATCTCACTCACCAATCA |
| chr19:11090988-11091987_567 | ACTCAGTTTTACAAAAGAAAGAGAGGCACTGGATTAATGTGTATCTCACTCACCAATCAACCTCTTCCTTAAGAGAAAAT |
| chr19:11090988-11091987_588 | AGAGGCACTGGATTAATGTGTATCTCACTCACCAATCAACCTCTTCCTTAAGAGAAAATGTTAAGGAAGTCTTAGGCAAG |
| chr19:11090988-11091987_609 | ATCTCACTCACCAATCAACCTCTTCCTTAAGAGAAAATGTTAAGGAAGTCTTAGGCAAGGCCTTGTTTGTTCATCACTTT |
| chr19:11090988-11091987_630 | CTTCCTTAAGAGAAAATGTTAAGGAAGTCTTAGGCAAGGCCTTGTTTGTTCATCACTTTAGTTTCTCTCTCCCGGGATGG |
| chr19:11090988-11091987_651 | AGGAAGTCTTAGGCAAGGCCTTGTTTGTTCATCACTTTAGTTTCTCTCTCCCGGGATGGCTGAGAATGTGATGTTTCCTC |
| chr19:11090988-11091987_672 | TGTTTGTTCATCACTTTAGTTTCTCTCTCCCGGGATGGCTGAGAATGTGATGTTTCCTCTGTTGTCAAGGAGACTACACC |
| chr19:11090988-11091987_693 | TCTCTCTCCCGGGATGGCTGAGAATGTGATGTTTCCTCTGTTGTCAAGGAGACTACACCCCTGATGTTTTCCTCCAGACT |
| chr19:11090988-11091987_714 | GAATGTGATGTTTCCTCTGTTGTCAAGGAGACTACACCCCTGATGTTTTCCTCCAGACTTCTGAGAGCTGGTGTGTGTTT |
| chr19:11090988-11091987_735 | GTCAAGGAGACTACACCCCTGATGTTTTCCTCCAGACTTCTGAGAGCTGGTGTGTGTTTCTAGCACTTTCTAGCTGCACC |
| chr19:11090988-11091987_756 | ATGTTTTCCTCCAGACTTCTGAGAGCTGGTGTGTGTTTCTAGCACTTTCTAGCTGCACCACCTCACGCTGTAGCTGGCTT |
| chr19:11090988-11091987_777 | AGAGCTGGTGTGTGTTTCTAGCACTTTCTAGCTGCACCACCTCACGCTGTAGCTGGCTTCAAGGCATATCCAGGGGGGAG |
| chr19:11090988-11091987_798 | CACTTTCTAGCTGCACCACCTCACGCTGTAGCTGGCTTCAAGGCATATCCAGGGGGGAGTTTCTTGTCCATTTCCTTTAC |
| chr19:11090988-11091987_819 | CACGCTGTAGCTGGCTTCAAGGCATATCCAGGGGGGAGTTTCTTGTCCATTTCCTTTACAAAGGGAAGTTGTTGGAATCT |
| chr19:11090988-11091987_840 | GCATATCCAGGGGGGAGTTTCTTGTCCATTTCCTTTACAAAGGGAAGTTGTTGGAATCTGAACCGCAAGCCTTCACTTAG |
| chr19:11090988-11091987_861 | TTGTCCATTTCCTTTACAAAGGGAAGTTGTTGGAATCTGAACCGCAAGCCTTCACTTAGACCAAAATCAGGCAACAGCGG |
| chr19:11090988-11091987_882 | GGAAGTTGTTGGAATCTGAACCGCAAGCCTTCACTTAGACCAAAATCAGGCAACAGCGGTGAGCGCAGCTCCAAACGTGT |

| | |
|---|---|
| chr19:11090988-11091987_903 | CGCAAGCCTTCACTTAGACCAAAATCAGGCAACAGCGGTGAGCGCAGCTCCAAACGTGTCAATGACTCACCCAAATTTGA |
| chr19:11090988-11091987_919 | GACCAAAATCAGGCAACAGCGGTGAGCGCAGCTCCAAACGTGTCAATGACTCACCCAAATTTGAGTAAGGGAGTTGGCTG |

**Supplemental Table 4: TaqMan Probes for qPCR/ddPCR.**

| **Target** | **Catalog number or Sequence** |
|---|---|
| *B2M* | #4326319E (Thermo Fisher Scientific Inc.) |
| *PPIA* | #4326316E (Thermo Fisher Scientific Inc.) |
| *LDLR* | #Hs01092524_m1 (Thermo Fisher Scientific Inc.) |
| *LDLR* paRNA | Probe: CAGGGCAGGCGACTTGATTTGTT<br>Forward Primer: CCCGTGAAGCTCTGATTCCC<br>Reverse Primer: TCTTCACCGGAGACCCAAAT |
| *TERC* | Probe: ATTCCCTGAGCTGTGGGACGTG<br>Forward Primer: AAGAGGAACGGAGCGAGTC<br>Reverse Primer: CACCAACAGGAAAGCGAACT |

**Supplemental Table 5: Information for publicly available datasets.**

| **Dataset** | **Source** | **Type** |
|---|---|---|
| H3K4me3 ChIP-seq (HepG2) | GSM8491771 | ChIP-seq |
| H3K27ac ChIP-seq (HepG2) | GSM8491772 | ChIP-seq |
| H3K4me1 ChIP-seq (HepG2) | ENCFF412ZOZ | ChIP-seq |
